# Structure determination of protein-peptide complexes from NMR chemical shift data using MELD

**DOI:** 10.1101/2021.12.31.474671

**Authors:** Arup Mondal, G.V.T. Swapna, Jingzhou Hao, LiChung Ma, Monica J. Roth, Gaetano T. Montelione, Alberto Perez

## Abstract

Intrinsically disordered regions of proteins often mediate important protein-protein interactions. However, the folding upon binding nature of many polypeptide-protein interactions limits the ability of modeling tools to predict structures of such complexes. To address this problem, we have taken a tandem approach combining NMR chemical shift data and molecular simulations to determine structures of peptide-protein complexes. Here, we demonstrate this approach for polypeptide complexes formed with the extraterminal (ET) domain of bromo and extraterminal domain (BET) proteins, which exhibit a high degree of binding plasticity. This system is particularly challenging as the binding process includes allosteric changes across the ET receptor upon binding, and the polypeptide binding partners can form different conformations (e.g., helices and hairpins) in the complex. In a blind study, the new approach successfully modeled bound-state conformations and binding poses, using only backbone chemical shift data, in excellent agreement with experimentally-determined structures. The approach also predicts relative binding affinities of different peptides. This hybrid MELD-NMR approach provides a powerful new tool for structural analysis of protein-polypeptide complexes in the low NMR information content regime, which can be used successfully for flexible systems where one polypeptide binding partner folds upon complex formation.

## INTRODUCTION

Molecular modeling has become an integral toolset for predicting bound conformations in structural biology. These successes are attributable to advances in protein structure prediction, robust docking pipelines for small molecules, and accurate free energy methods for quantifying relative (and absolute) binding affinities^1^. Despite these advances, the accuracy of these methods decreases rapidly for systems involving significant conformational changes upon complex formation, where receptors can accommodate multiple binding modes, and for highly charged systems^2,3^. In particular, systems involving disorder-to-order transitions upon complex formation, including peptides that fold as they bind, challenge current protein-peptide docking methods. Recently, machine learning tools have brought fast and accurate predictions for protein structures^4,5^ and are now being extended to predictions of complexes^5,6^. More generally, three-dimensional structures of peptide-protein complexes provide important information for understanding the mechanisms of multiprotein complex assembly, and have the potential to inform drug discovery.

Here we describe an integrative approach to structure determination for peptide-protein complexes combining NMR chemical shift data and molecular simulations. High information-content NMR studies rely on extracting many distance and orientation restraints to solve the structure of the peptide-protein complex^7^. At the other extreme, lower information-content NMR studies, such as backbone chemical shift data which is prerequisite to more extensive studies, provide valuable information about the binding epitope and (in some cases) the bound-state conformation of the peptide, but do not usually provide enough data to reliably characterize the binding mode and structure of the resulting complex.

Molecular simulations approach the problem of peptide binding by sampling the binding/unbinding landscape, including multiple binding modes and peptide conformations, relying on statistical mechanics to identify preferred conformations in the ensemble. Sampling multiple binding/unbinding events requires timescales much longer than the bound-state lifetime^8^, entailing a large computational effort even with special purpose computers^9^ or advanced sampling technique strategies^10–14^. Integrating experimental data reduces the conformational space, focusing sampling on structures that satisfy the physics as well as the experimental data^15,16^. In this work we identify synergies between incomplete or “sparse” NMR data and simulations for structure prediction of peptide-protein complexes, focusing on applicability and transferability as well as limitations. This pipeline allows more rapid structure determination of complexes than conventional NMR approaches, and can provide structures of complexes even for systems for which extensive NMR data cannot be obtained. We focus on the binding of polypeptides to the extra-terminal domain (ET) of bromo and extra-terminal domain (BET) proteins, which exhibit disorder to order transitions of the polypeptide upon binding, allosteric changes in the receptor, and accommodate peptides binding in different conformations^7^. This biologically-important system exhibits a large degree of plasticity in binding modes and peptide conformations ^7,17–19^,, as well as a large range in binding affinities, and poses challenges to current computational and experimental approaches.

The BET family of proteins (BRD2, BRD3 and BRD4) play important roles in eukaryotic gene regulation by recognizing and binding epigenetic signatures and recruiting other regulatory proteins. Structurally, BET proteins contain two bromodomains that recognize and bind acetylated signatures in histones, and an extra-terminal domain (ET) which serves as an anchor point to recruit other proteins such as NSD3, JMJD6, CHD4, GLTSCR1 and ATAD5^19^. Some viruses like the murine leukemia virus (MLV) or Kaposi’s sarcoma-associated herpesvirus (KSHV), encode proteins that also bind to the ET domain. For retroviral integration, this effectively facilitates their location near active transcription start sites and CpG islands^20^. The MLV integrase (IN) contains an intrinsically disordered C-terminal polypeptide “tail” segment that becomes structured upon binding the ET domain. KSVH virus has a latency-associated nuclear antigen (LANA) protein that also binds ET. Understanding and predicting how different polypeptide sequences bind the ET domain can potentially lead to new approaches for cancer treatment and gene replacement therapy.

The ET domain is an 87-residue three-helix bundle, with a binding site defined by a hydrophobic pocket flanked by a negatively charged loop region connecting helices α2 and α3. Proteins interact with the ET domain through short peptide epitopes which anchor hydrophobic residues in ET’s hydrophobic pocket and interlace positively charged residues of the ET-binding polypeptide segment with negatively charged residues of ET through a zipping mechanism^17^. Interestingly, the binding mode, orientation of the bound polypeptide segment, and even secondary structure of the bound polypeptide can change considerably for different polypeptide sequences, causing the loop in the ET receptor to adopt different conformations in different ET-polypeptide complexes (see Fig. 1).

**Figure 1.**
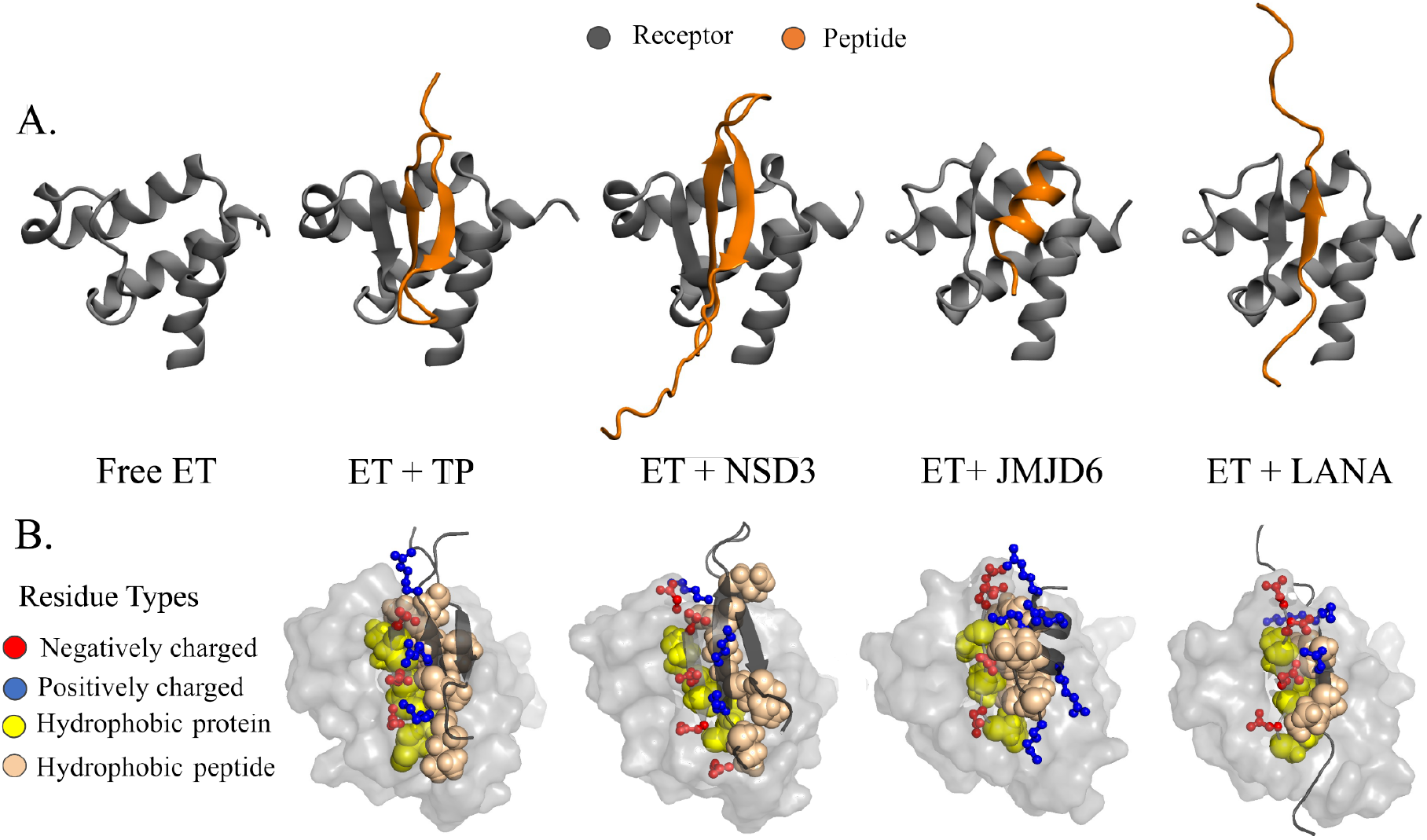
Plasticity of ET - peptide complex formation. (A) Experimental structures of the ET domain of BET proteins interacting with different peptides highlight differences across the binding modes observed in different ET – peptide complexes. (B) A network of alternating positively (blue) and negatively (red) charged interactions between the peptide and protein residues a zipper like interacting mechanism which is further stabilized by hydrophobic packing between a cleft in the protein and hydrophobic peptide sidechains.

We formulated this study in two stages. First, we carried out blind computational modeling of two peptide-protein complex structures, followed by assessment of model accuracy based on high-quality NMR structures of these complexes^7^. Here, the experimental team provided NMR datasets for peptide-protein complexes for which structures were not yet deposited in the Protein Data Bank, and not available to the prediction team, with increasing information content, and collected predictions from the computational group at each stage. The lowest tier data used only backbone ^15^N-^1^H chemical shift perturbation (CSP) data measured on the receptor protein – effectively identifying possible binding hot-spots. For ET these CSP data include both effects at the peptide binding site, and changes throughout the structure due to allosteric conformational shifts that result from peptide binding^7^. At the highest tier of experimental information, in addition to the backbone ^15^N-^1^H CSPs measured on the receptor protein, dihedral angle restraints based on backbone chemical shift data for the isotope-enriched receptor-bound peptide^21,22^, and a few peptide-protein contacts based on the strongest NOEs observed between the protein and bound peptide, were provided. In the second stage of the study, we also extended the method to other peptide - ET domain complexes for which three-dimensional structures were already published, to assess the generality of our methods.

## METHODS

### Experimental Methods

#### NMR Data

Experimental NMR data were generated for murine BRD3 ET domain (residues 554 to 640) and for its complexes with the 23-residue C-terminal tail peptide (TP) of MLV IN (residues 1716 to 1738 of the Gag-Pol polyprotein), and the ET-binding polypeptide segments of murine NSD3 (residues 151 to 184), as summarized in Table 1. ^13^C,^15^N-enriched samples of ET were produced using standard methods^7,23^, while isotope-enriched peptides were produced as fusion proteins followed by proteolytic cleavage, as described elsewhere^7^. Sequence-specific resonance assignments for peptide complexes (for both the ET protein and for the bound polypeptides) were determined using standard triple-resonance NMR methods, also described elsewhere^7^. NMR data for ET-binding domains of CHD4^24^ (BMRB ID 30367), BRG1^24^ (BMRB ID 30368), LANA^17^ (BMRB ID 26042) and JMJD6^18^ (BMRB ID 30373) were obtained from the BioMagResDataBank (BMRB). Backbone amide ^15^N-^1^H chemical shift perturbations (CSPs) of apo BRD3 ET domain relative to values in the complex were calculated using Δδ_(N,H)_ = ((Δδ_N_/6)^2^+(Δδ_H_)^2^)^0.5^, and plotted as a function of BRD3 ET sequence. The threshold for defining a significant CSP was determined by iterative analysis^25^. The standard deviation (σ) of the shift changes Δδ_(N,H_ was first calculated. To prevent biasing the distribution by including the small number of residues with very large shift changes, any residues for which the shift change is greater than 3σ were excluded. The standard deviation (σ) of the remaining Δδ_(N,H)_ values was then recalculated. Iteration of these calculations was performed until no further residues were excluded. The threshold value for a significant CSP was then set to Δδ_(N,H)_ = 3σ = 0.02 ppm. ^13^C and ^15^N-edited 3D NOESY spectra for uniformly ^13^C,^15^N-enriched ET-TP and ET-NSD3 complexes were recorded with NOE mixing times of 120 ms.

**Table 1.**
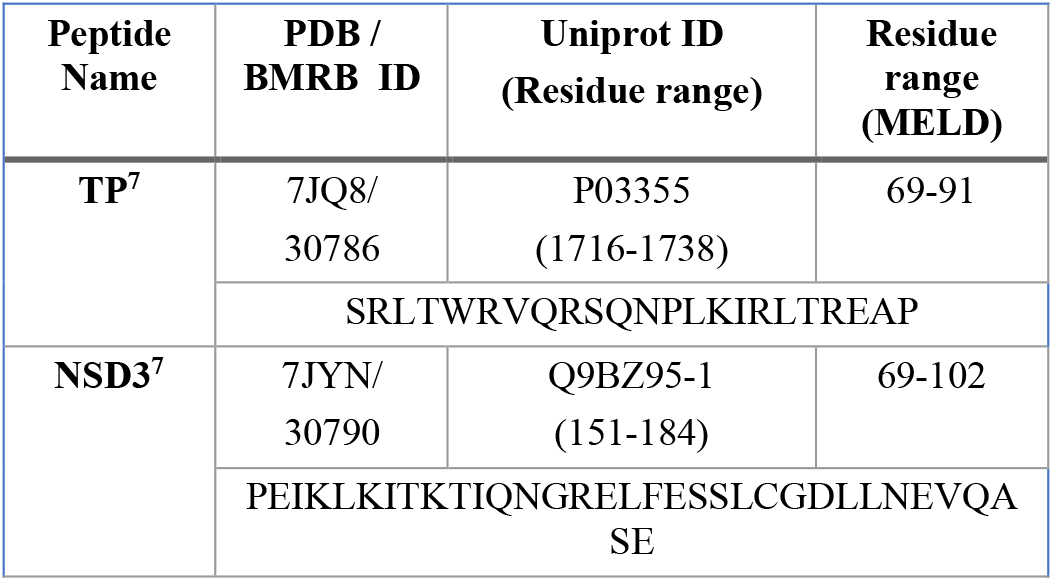
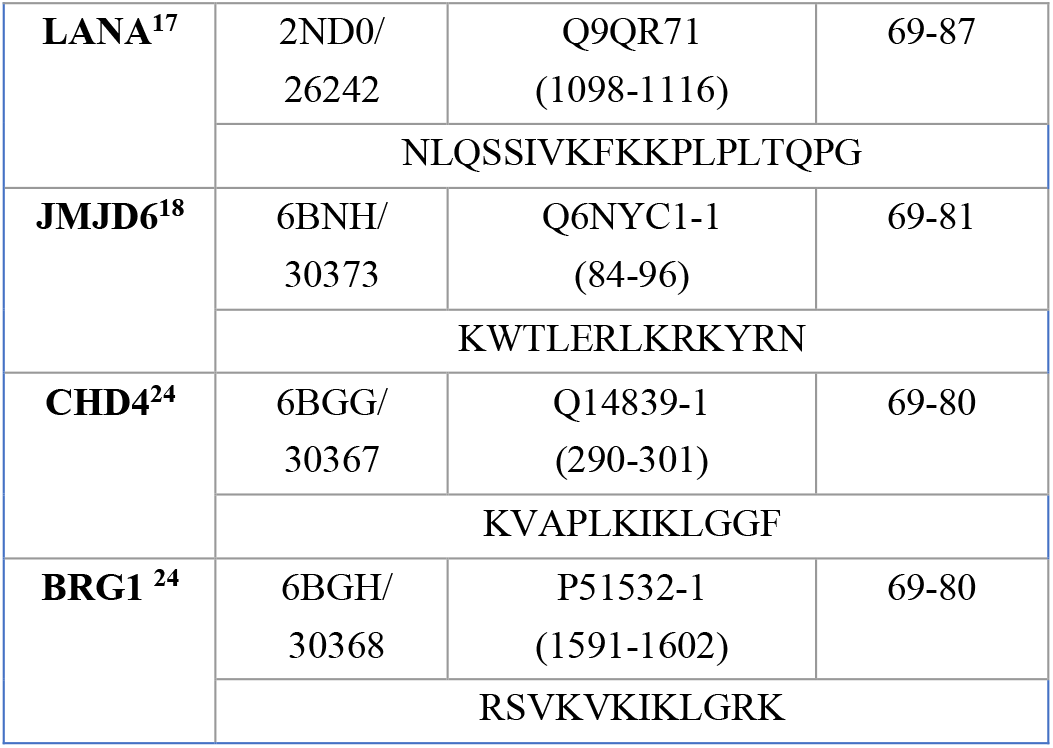
Six systems investigated with MELD-NMR and their corresponding peptide sequences and public database identifiers, where peptide residue numbers correspond to the sequence numbering used for these studies.

#### Isothermal Titration Calorimetry

Isothermal titration calorimetry was carried out using a MicroCal VP-ITC Isothermal Titration Calorimeter located in the Analytical Biochemistry Core Facility of the Center for Biotechnology and Interdisciplinary Sciences (CBIS) at Rensselaer Polytechnic Institute. Recombinant TP (SRLTWRVQRSQNPLKIRLTREAP) and NSD3 (EFTGSPEIKLKITKTIQNGRELFESSLCGDLLNEVQASE) were prepared as described previously^7^. Samples of ET(∼ 2.4 ml) and peptide binding partners (∼ 300 μL) were prepared for ITC studies by dialyzing together in separate dialysis bags placed the same beaker of the ITC buffer containing 25 mM Tris, 100 mM NaCl, 5 mM TCEP at pH 7.5. The ET and peptide binding partners were first dialyzed in 1L ITC buffer at 4 °*C* for 8 hours, and then dialyzed into a new 1L ITC buffer at 4 °*C* overnight. Protein and peptide concentrations were determined after dialysis by absorbance spectroscopy at 280 nm or 205 nm using extinction coefficients for ET (e_280_ = 4470 M^- 1^cm^-1^), TP (ε_280_ = 5500 M^-1^cm^-1^) and NSD3 (e_205_ = 126,480 M^- 1^cm^-1^, contains no Tyr or Trp), calculated from their respective amino acid sequences.

#### Datasets used for the blind study

We use three different experimental NMR datasets to mimic different approaches: (1) **CSP**, (2) **CSP+TALOS**, (3) **CSP+TALOS+NOE**. For the lowest information content dataset, experimental docking data include only backbone ^15^N-^1^H CSP data for the protein receptor in the presence/absence of a bound peptide. We call these the **CSP datasets**. This approach uses comparison of the [^15^N-^1^H]-HSQC spectra of apo and peptide-bound receptor, and has the advantage that the peptide binding partner does not require isotope enrichment. Residues of the receptor (in this case, ET) for which there is a chemical shift perturbation upon complex formation may be directly involved in binding or indirectly affected by allosteric conformational changes. Hence, these CSP data do not provide information about which specific atoms of the protein (e.g. backbone vs. sidechain) are involved in the interaction with the peptide, or which peptide residues might be involved in the binding. The CSP threshold for significant perturbations (calculated as described above) indicates that the ^15^N-^1^H chemical shifts of the majority of residues in the ET receptor are perturbed upon complex formation – with a broad range of CSP values (Fig. S1). Based on the distribution of CSPs across the ET structure, for both the ET-TP and ET-NSD3 complexes we assigned only residues with relatively large CSP values > 0.25 ppm as “active residues”, likely to be in or near the peptide binding site. These include 20 and 22 protein residues, respectively, for the two complexes (Fig. S1).

The second dataset adds conformational restraints for phi/psi dihedral angle ranges for the bound peptide based on the backbone chemical shifts measured for the bound peptide in the complex, determined with the program TALOS^21,22^ (**CSP+TALOS** datasets). This data requires isotope-enrichment of the peptide ligand, and NMR assignments for the bound peptide. TALOS-generated peptide dihedral torsion restraints were used in MELD simulation with maximum and minimum values shown in SI Tables S1 and S2. The third dataset includes restraints derived from three strong interchain NOEs among backbone and relatively easy to assign sidechain resonances (**CSP+TALOS+NOE** datasets). Specifically, in this third data set distance restraints were imposed between: Val 24 CG1 / Trp 73 HH2, Ile 44 CD1 / Ile 84 CD1, and Glu 47 CG / Leu 82 CD2 for the ET-TP system, and between Ile 44 H / Leu 73 H, Ile 42 H / Ile 75 H, and Val 24 CG1 / Phe 86 CD1 for the ET-NSD3 system. Simulations were performed by trimming off 19 unstructured residues in the N-terminal region of ET (which are distant from the peptide binding site) and renumbering the resulting trimmed domain sequence to start at residue 1. Thus, the residue numbering convention used in the MELD-NMR calculations uses residues 1 to 68 for the ET receptor (corresponding to residues 573 to 640 of the BRD3 protein), with residues of the peptide binding partner numbered from 69 onward (corresponding residue numbers shown in Table 1).

### Computational methods

#### MELD approach

MELD is a plugin to the OpenMM^26^ molecular simulations engine. It integrates data and simulations through Bayesian inference^15^. The main advantage is that it accommodates data sources that are ambiguous and/or noisy, making it well-suited for analysis of protein NMR data.. For example, CSP information on the ET receptor protein identifies possible sites of interaction between the protein and peptide – but we do not know which residues in the interface contact with residues in the bound peptide, and which are never in direct contact (e.g., CSPs distant from the binding site can arise from allosteric or propagated structural changes). We thus include all possibilities, knowing that only a small subset will be present in the structure of the complex (see Fig. 2). MELD samples through conformations and different implementations (subsets) of the data, to produce an ensemble that is compatible with a subset of data and the physics model (given a force field).

**Figure 2.**
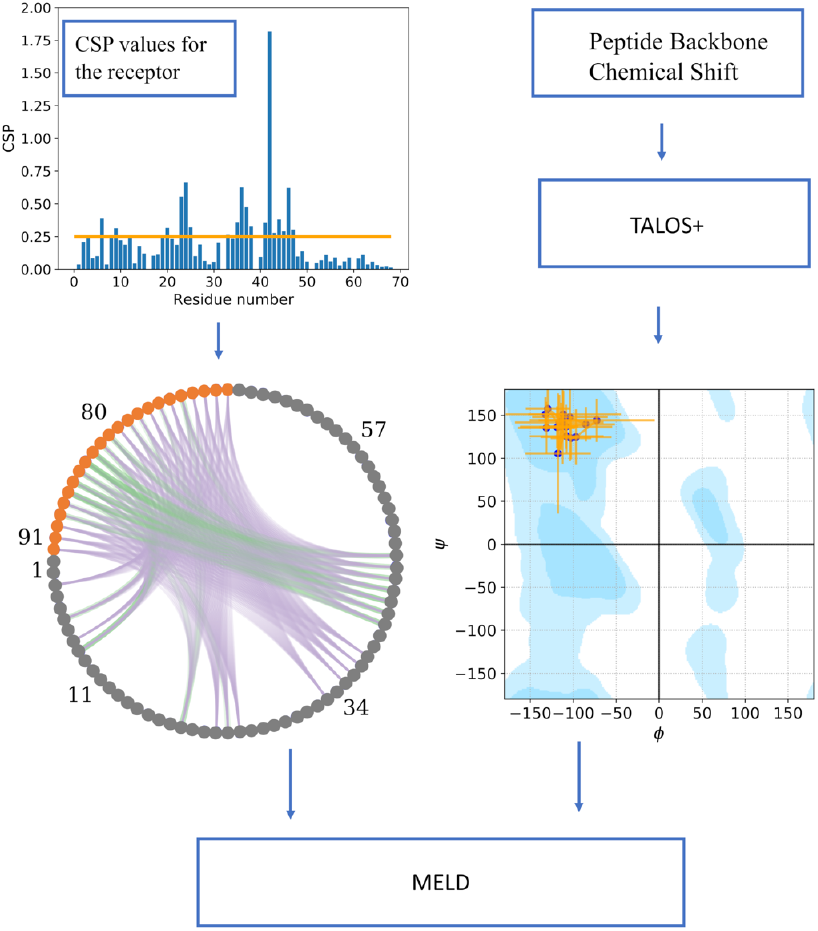
General flowchart of the current work. The upper left plot shows NMR CSP values for each residue in the receptor, with the threshold cutoff used to define active residues shown as an orange line. Given the active residues, the lower left circular plot represents the combinatorics of possible restraints between the peptide (orange circles) and the receptor (grey circles) – each line between pairs of residues is a potential contact. Purple lines represent contacts that are not present in the native structure, and green lines shows the ones present (true positives). For some active residues there are no green lines, indicating that their high CSP values were due to an allosteric or propagated structural-changes. The lower right plot shows the dihedral restraints on the binding peptide conformation, with their uncertainties (orange lines) used in MELD with TALOS data. The Ramachandran plot is made using https://github.com/gerdos/PyRAMA.

#### MELD simulations

As in previous studies, we used an H,T-REMD protocol^27^ for efficient exploration of the energy landscape. We used 30 replicas running for 1.5 µs starting from an unbound folded receptor and extended peptide conformation placed 30 Å away from the receptor. The temperature increases geometrically from 300 K at lowest replica to 500 K at the 12^th^ replica and is kept constant at 500 K afterward. The Hamiltonian changes according to how strongly we enforce the data (strongly, k=350 kJ/(mol nm^2^), below replica 12^th^, and with no restraints at the 30^th^ replica—changing non-linearly in between the 12^th^ and 30^th^ replicas). The physics model uses the GBNeck2 implicit solvent model^28^ and a combination of the ff14SB^29^ (side chains) and ff99SB^30^ (backbone) force fields.

The CSP data was modeled by all possible combinations between the set of all peptide residues and the set of ET *active* residues with CSP above the Δδ_(N,H)_ = 0.25 ppm threshold, using a flat-bottom harmonic restraint between C_β_’s of each pair. The restraints added no energy penalty up to 8 Å, and then the energy penalty increased quadratically until 10 Å, and linearly beyond, with a force constant of 350 kJ/(mol nm^2^). The combinatorics leads to many possible restraints, but only a few are present in the bound structure – hence we assigned a 4% confidence level to this dataset. Which restraints are enforced is deterministic: at each timestep all restraint energies are evaluated, and the *4%* lowest energy restraints are enforced until the next timestep. Different replicas can satisfy different sets of restraints. An advantage of MELD is that each different data source can have different confidence values. For example, a second protocol created combinatoric restraints between only the hydrophobic residues in the peptide and the *active* protein residues. This resulted in a lower number of overall restraints and a higher confidence level (10%).

We modeled bound-peptide chemical shift data in a similar manner by enforcing *phi* and *psi* dihedral angle restraints based on the minimum and maximum values provided by TALOS (see Fig. 2 and Tables S3-S6). Modeling of dihedral restraints is based on TALOS analysis of the chemical shift values for each residue, and are therefore of higher accuracy than the ambiguous modeling of CSP data. We set the confidence on this data to 80%. Finally, we modeled NOE data with a 5 Å flat-bottom potential between backbone-bacbkone hydrogens. For NOEs between sidechain-sidechain or backbone-sidechain hydrogens, we mapped the sidechain hydrogen to the corresponding heavy atom and added a 6 Å fat-bottom potential. We used 100% confidence on this dataset.

In all simulations, we also applied internal distance restraints to the ET structure in order avoid protein unfolding at high temperatures in the replica exchange. For this purpose, we calculated all C_α_ -C_α_ distances in the apo-protein, selected those closer than 8 Å, and created a restraining potential for each using an 8 Å flat-bottom harmonic potential. To allow for possible conformational changes during binding, both locally to the binding site due to binding plasticity and distant from the binding site due to allosteric changes, we set this dataset with a 90% confidence inside MELD. The accuracy parameter in the different data sources plays a critical role in guiding the search in MELD – but, it is unknown *a priori*. Enforcing higher accuracy values result in more restrained systems and thus faster convergence (shorter simulation times). However, if the accuracy value is set too high, the restrained protein structure might not be compatible with all of the data, leading to incorrect simulations.

#### Competitive binding study

Relative binding affinities can be calculated from MELD simulations in which two (or more) peptides compete for the binding site. The relative binding affinities for ET was assessed for the TP and NSD3 peptide sequences shown in Table 1. We selected 3 possible restraints common to both systems based on the experimentally-determined structures of these complexes. In the starting conformation both peptides where 30 Å away from each other and from the receptor. In this case the contacts we enforced had to be satisfied by either peptide at lower replicas, whereas both were unbound at higher replicas. We also added restraints to keep the two peptides from interacting with each other. The ratio of the populations of peptide molecules bound in the binding site is related to the relative binding free energy^31^.

#### Clustering

We cluster the last 500 ns of the five lowest temperature replicas using RMSD as a similarity metric with a hierarchical agglomerative clustering algorithm implemented in *cpptraj*^32^. All protocols for complexes use average linkage to calculate distance between clusters with a 1.5 Å cutoff. For all cases, we have two clustering protocols: one using LRMSD (Ligand root mean square deviation)^33^ calculated considering the whole peptide and another using LRMSD calculated on only the core region of the peptide (excluding floppy terminal regions). We report the centroid of the highest populated cluster as our prediction.

#### AlphaFold predictions

We used the AlphaFold^4^ advanced colab version (https://github.com/sokrypton/ColabFold)^34^ to predict structures of these 6 complexes. For predicting protein-peptide complexes, a 30-residue glycine linker was used between the C-terminal end of the protein and the N-terminal end of the peptide sequence, to create a single polypeptide chain that was used as input to AlphaFold. Five models were predicted for each complex, and ranked according to their pLDDT scores^4^. A similar protocol has recently been reported to provide accurate results in predicting other protein-peptide complexes^35^. Related ET-peptide structures already published in the PDB were excluded for AlphaFold analysis. Each complex model was then refined with the ff99SB forcefield.

#### Analysis methods

Predicted structures were assessed using different state-of-the-art metrics used in the protein-protein or protein-peptide docking communities including IRMSD, f_nat_ and IRMSD^2,33,36^. Experimental structures determined by NMR methods were reported as ensembles with 20 models. For calculating each of these metrics, the medoid structure of each NMR ensemble was used. The medoid, or representative, structure for each complex was calculated using the *PDBstat* software package^7,37^.

## RESULTS

### Isothermal Titration calorimetry measurements

Binding affinities of TP and NSD3 peptides to BRD3 ET, K_d_ ∼ 100 nM and ∼ 10 μM, respectively, were estimated at pH 7.5 and 25 °C by preliminary isothermal titration calorimetry measurements. The former value is similar to the reported K_d_ ∼ 150 nm for TP – BRD4 ET binding at pH 7.0 and 25 °C^38^ ; there is an ∼100-fold difference in affinities for ET between TP and NSD3 peptides.

### Blind studies distinguish different binding modes

Figure 3 highlights our results using different datasets for the two blind studies (TP and NSD3 peptides binding to the BRD3 ET domain). In all cases we report the centroid of the top population cluster from the ensembles. Interestingly, each peptide requires a different amount of data to accurately determine a complex structure. MELD-NMR using CSP for the ET receptor and TALOS backbone dihedral-angle restraints for the bound peptide (i.e. only chemical shift data) successfully identifies both peptides binding through antiparallel strands, with TP forming intermolecular antiparallel beta-sheet interactions with ET along its C-terminal region, and NSD3 forming intermolecular beta-sheet interactions through its N-terminal region, resulting in flipped orientations for the peptide hairpin with respect each other (see Fig. 3). These binding modes are in excellent agreement with the experimental NMR structures of the two complexes (IRMSD = 2.28 Å and 1.97 Å, respectively). For the stronger-binding peptide (TP), CSP on ET data alone was sufficient to provide an accurate binding mode (see Fig. 3), and no data for the bound peptide was needed. Using the CSP on ET data from TP for predicting the NSD3 binding mode did not change the predictions (see SI), demonstrating that even with a significantly-different binding mode, a single study of CSPs on the receptor can be used to successfully guide binding of other binding peptides. For both complexes, adding the three strongest intermolecular NOEs does not increase the accuracy of the prediction.

**Figure 3.**
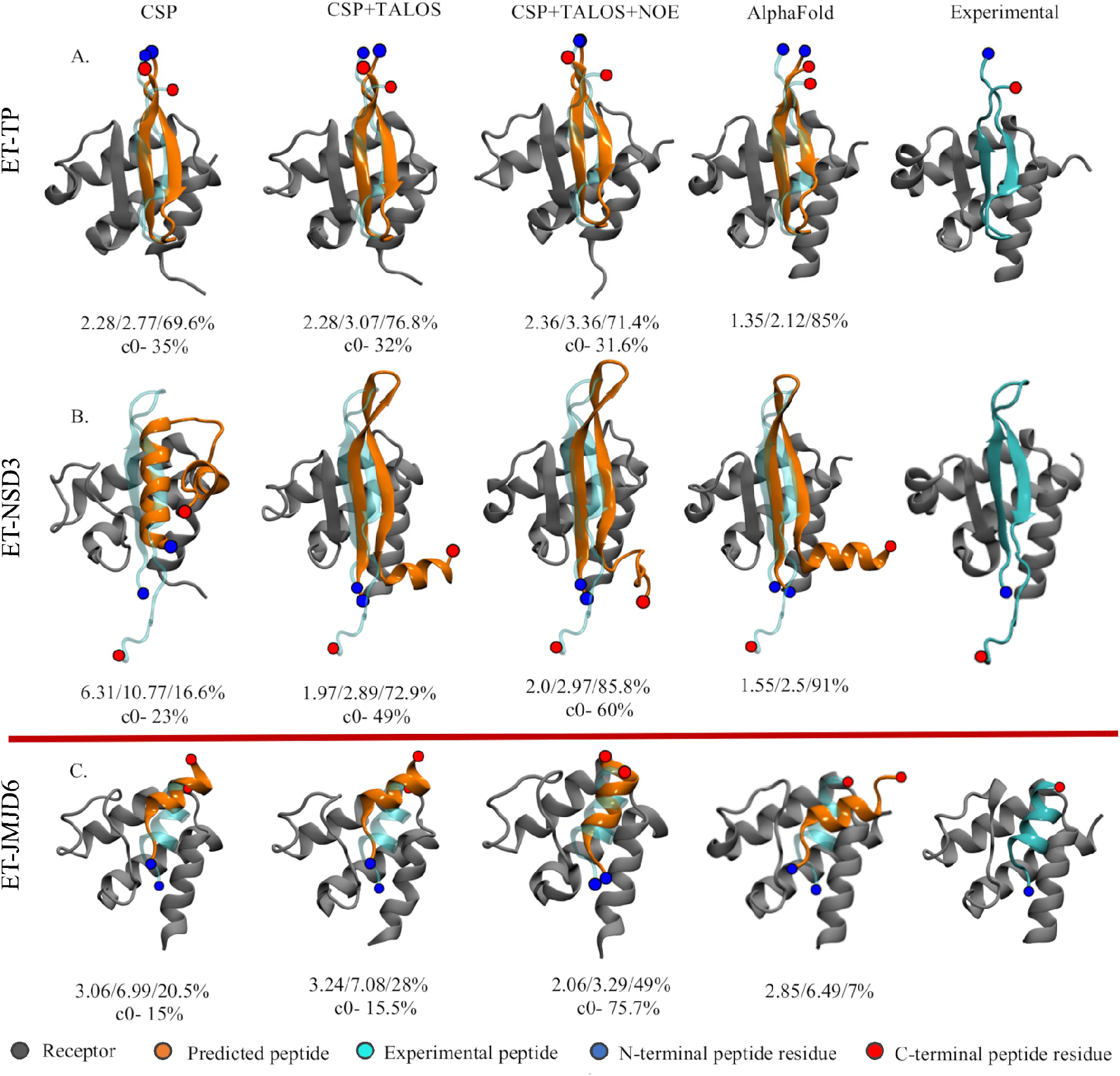
Predicted binding modes. The top panel shows the top MELD prediction using different experimental datasets (first three columns) for the blind study along with AlphaFold (4^th^ column) predictions: (A.) BRD3-TP, and (B.) BRD3-NSD3. Panel C shows the predictions for the known helical binder, BRD3-JMJD6. The numbers below each structure represent IRMSD/ILRMSD/f_nat_ in the first row and the population of the top clusters in the second row.

Interestingly, using no experimental data, for these two complexes AlphaFold performs similarly to chemical-shift guided MELD (IRMSD = 1.35 Å and 1.55 Å, respectively). For the ET-NSD3 complex, AlphaFold also predicts an additional alpha helix in the tail of NSD3, which is unstructured in the experimental NMR structure (see Fig. 3 and Fig. S2). These results demonstrate the potential to use AlphaFold either to screen for potential binding poses prior to beginning experimental studies, such as NMR-guided MELD modeling, or for providing a validation of the results of MELD+NMR studies.

Funneling plots capture the ability of MELD ensembles to sample and direct towards the native complex. Each funneling plot shows all the cluster centers in the ensemble as a function of the RMSD to the complex structure (see Fig. 4). MELD simulates multiple binding/unbinding events driven by the data. Unbound states are represented as low population clusters sampled at high replica index (red). At lower replica indexes, the simulation samples bound (green clusters) and misbound (blue clusters) states. We seek approaches that have funneling towards the green regions – as seen for the TP peptide. Although the final top representative structure is identified based on clustering on the lower replicas, these funneling plots clustering on all replicas are useful to identify confidence in the results of MELD +NMR. When the experimental structure is unknown (e.g. during these blind studies) we use the top scoring cluster as a reference for RMSD calculations. We used these types of plots to determine the CSP thresholds and determine the *accuracy* parameter as a self-consistency test before submitting predictions. For the TP peptide different protocols agreed on the same bound conformation, increasing the confidence in our predictions (see Fig. S3). For the NSD3 peptide, using CSP data alone, different protocols were not in agreement (see the higher number of misfolded states in Fig. 4), with resulting conformations differing by more than 5 Å backbone RMSD between them. Adding backbone dihedral restraints for the bound peptide determined from chemical shift data using TALOS narrowed the number of clusters, and helped identify a core region that was bound, with a flexible terminal region. This flexible region was responsible for a higher diversity of binding modes and lower populations of the top cluster – clustering on the core region rapidly identified a top cluster with highest confidence (49% population of the top cluster). *A posteriori* analysis looking back at the CSP only dataset shows that the correct binding mode was identified as the 5^th^ cluster – what in the docking field is considered a s*coring failure*. Adding dihedral restraints for the bound peptide reduces the number of clusters by 50% with respect to the initial protocol (see Fig. S3). Similarly, *a posteriori* analysis for TP reveals that excluding the disordered terminal region residues from the clustering calculation improved the confidence score for the correct prediction (populations above 55%, see Fig. S3).

**Figure 4.**
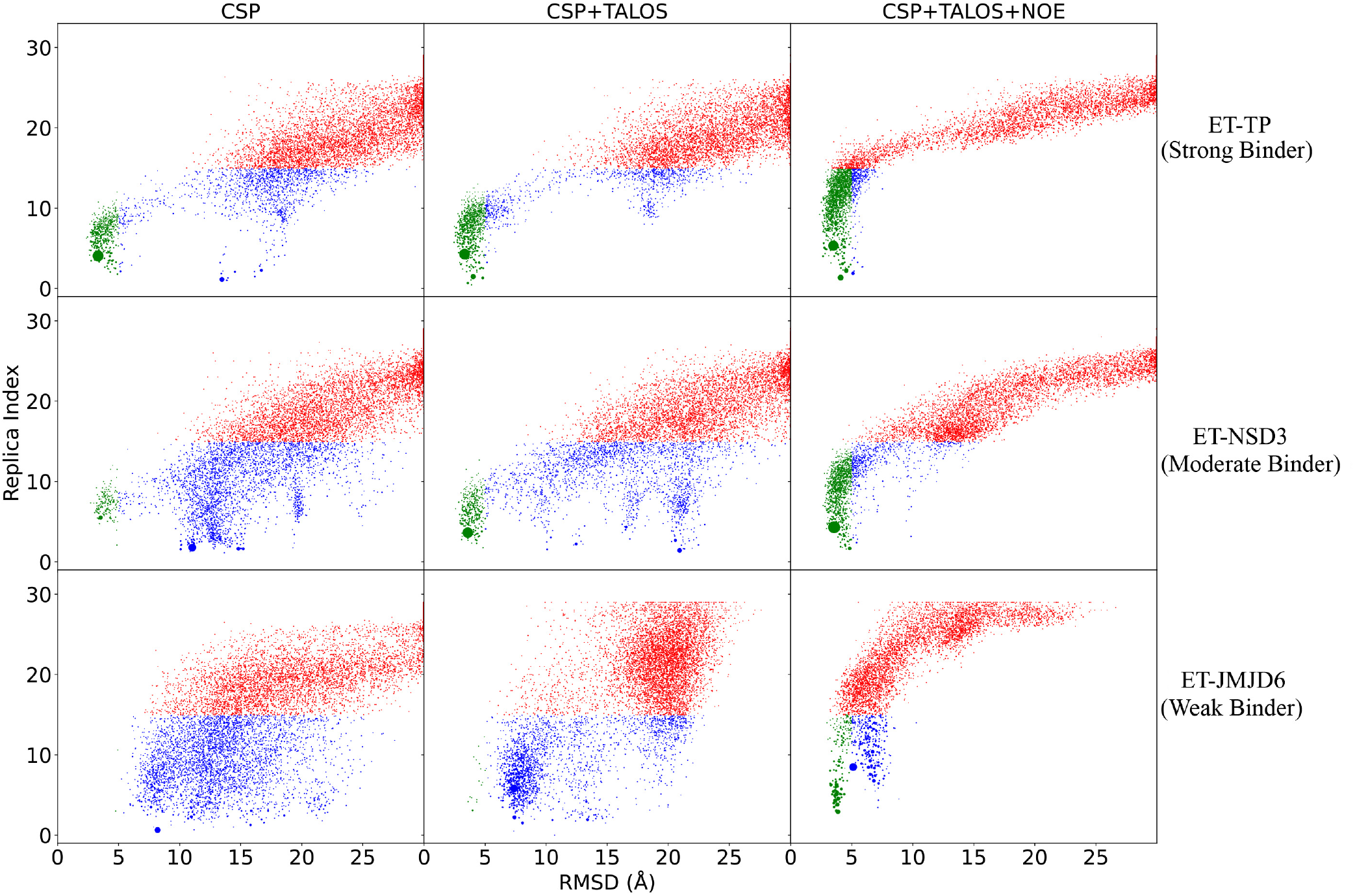
Binding funnels plot. Each point represents the centroid of a cluster center arising from the clustering of MELD ensembles. The size of the cluster is proportional to the population. Each cluster centroid is represented at the average replica index of the cluster to which they belong and the LRMSD of the centroid structure with respect to the experimental one. Clusters sampled at high replica index are shown in red, clusters sampled at lower replicas are shown in green (LRMSD < 5Å) or blue. Blue represent mis-bound conformations. Successful simulations have high populations of green clusters.

### Studies on known peptide-ET complexes

Four additional peptide-ET complexes have been previously experimentally characterized (JMJD6, LANA, CHD4 and BRG1). These four peptides are all weaker binders than TP (K_d_ ∼ 160 µM^18^, ∼ 635 µM^17^, ∼ 95 µM^24^, and ∼ 7 µM^24^ respectively). Of these, JMJD6 binds as an alpha-helix, with the rest binding as single anti-parallel β-strands. MELD-NMR calculations were carried out for the published complexes to further explore the interplay between binding affinity, information content of the data, and MELD’s ability to predict structures of even weaker-binding complexes such as JMJD6, LANA, and CHD4. JMJD6 is the only solved structure binding as a helix^18^, illustrating the broad binding mode plasticity of the ET domain. However, JMJD6 is also a special case, as access to the ET-binding epitope (amino-acid residues 84 to 96)^18^ requires a conformational change in the JMJD6 protein structure in order to expose these residues for binding into the ET cleft.

Lower affinity complexes have lower occupancy of the bound-state conformation, resulting in weaker CSP effects on the ET receptor (see Fig. S1). This challenge was addressed, assuming competitive binding of these peptides with TP, by using the CSP data for the ET-TP complex in MELD+NMR modeling of these other complexes. Similarly, fast or intermediate exchange on the NMR chemical shift timescale also results in ensemble-averaging of the bound-state peptide chemical shift data, precluding straight-forward determination of the bound-state peptide backbone dihedral angles from these chemical shift data for weakly-binding complexes using TALOS (see Tables S7-8). In principle, in the intermediate or fast exchange regime of these weaker complexes, bound-state peptide backbone chemical shifts can be determined using more sophisticated NMR experiments such as relaxation dispersion (e.g. Carr-Purcell-Meiboom-Gill^39^, CPMG), chemical exchange by saturation transfer (CEST)^40^ and/or peptide titration experiments. To mimic such NMR data for testing MELD-NMR’s applicability even for weakly binding systems, we obtained phi/psi restraint ranges directly from the experimental structures using *MDTraj*^41^. For consistency with the blind study, we simulated three strong NOEs based on the experimental structures (see Table S9) for generating intermolecular distance restraints for the CSP+TALOS+NOE dataset.

Of the four peptides, MELD recovers the experimental binding modes for tighter-binding CHD4 and BRG1 complexes using only CSP on ET data (see SI and Fig. S4). LANA is the weakest binding peptide in the set, and using CSP on ET data alone produces incorrect binding modes and peptide conformations. Adding peptide backbone dihedral angle restraints for the LANA-ET complex is enough to allow MELD+NMR to model a binding mode close to the experimental structure. Hence, for these three complexes, MELD-NMR provided accurate models using only chemical shift data. The accuracy of the LANA-ET binding pose is further improved by the addition of a few simulated NOE restraints (backbone-backbone contacts, see SI and Fig. S4).

The ET-JMJD6 complex, however, was more challenging. JMJD6 was found to bind as a helix using each of the three different datasets, with a bound-state structure similar to that observed in the experimental structure of this complex^18^. However, the helix is displaced from the experimental binding mode when using CSP and CSP+TALOS data. Interestingly, for this system AlphaFold also predicts a helical structure for the bound peptide. However, this binding pose is also significantly different from the reported experimental structure (Fig. S2). The top five AlphaFold predictions are diverse in terms of binding mode and more closely agree with the MELD predictions than with the experimental JMJD6-ET complex structure. As expected, adding three strong-NOE-based distance restraints simulated from the experimental structure to MELD+NMR simulations recovers the experimentally-observed binding pose (see Fig. 4).

### Competitive binding simulations identify TP as a stronger binder than NSD3, consistent with experimental ITC measurements

Although AlphaFold was able to predict structures of TP - ET and NSD3 - ET complexes quite well without any experimental data, this modeling approach does not provide any information about their binding affinities. However, MELD calculations include a complete statistical-mechanical energy function, and have the potential to also assess alternative predicted binding modes and their relative binding affinities, or the relative affinities of two binding peptides. For comparing the binding affinities of the TP and NSD3 peptides for the ET domain, we chose a common set of information to guide each peptide to the binding site (see Methods and Fig. 5A), and performed competitive binding simulations. Figure 5B summarizes the population of each peptide in the binding site according to replica. While at high replica index both peptides are unbound, early in the binding process there is a marked preference for TP binding over NSD3. The latter peptide can sample the binding site multiple times, especially in intermediate replicas but it is rarely sampled at the lowest replica. Based on these results we predict a ΔΔG_bind_ of -2.4 kcal/mol. The ITC experiments at pH 7.5 confirmed that the TP peptide is a better binder, with experimental relative binding free energy of about 2.6 kcal/mol.

**Figure 5.**
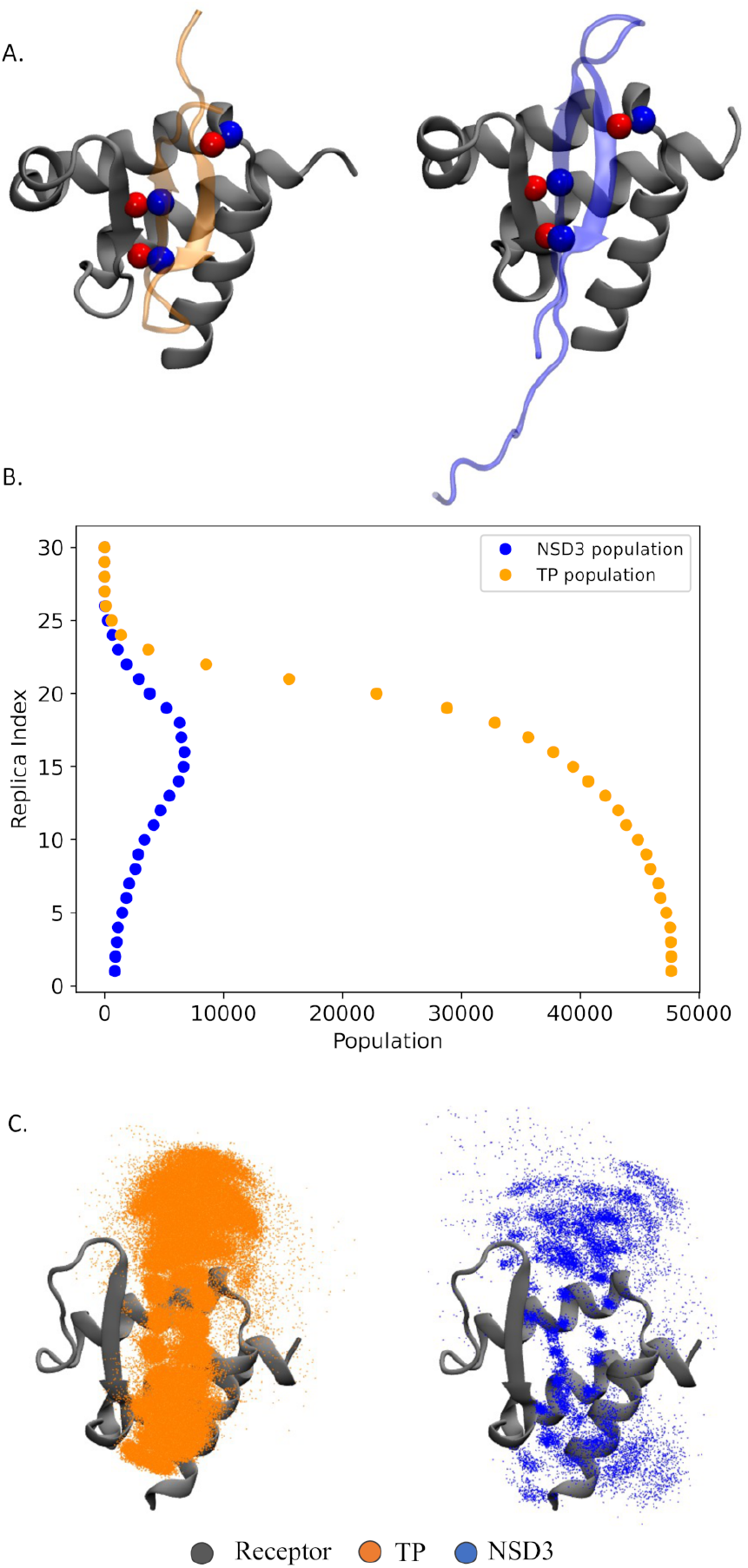
Competitive binding simulations. (A) Three equivalent handpicked contacts used to guide sampling for competitive binding. Red and Blue spheres represent Oxygen and Nitrogen respectively. (B) Population of native bound conformation for each peptide at each of the 30 replicas. (C) Superposition of bound peptides in the lowest temperature ensemble: each dot represents a C_α_ of the bound peptide.

## DISCUSSION

The protein-protein docking field has significantly advanced thanks to community efforts such as CAPRI (Critical Assessment of Prediction of Interaction)^42,43^. However, it is an unsolved challenge to reliably predict, without any experimental data, binding poses when a large degree of conformational flexibility is involved. It is particularly challenging to model bound peptide conformations in systems like these ET – peptide complexes involving disorder to order transitions upon complex formation. Such docking studies are particularly challenging for binding receptors like ET that exhibit binding plasticity, where peptides can bind along different modes and conformations. Moreover, while docking can be more successful when the bound-state peptide conformation is known, such docking studies do not provide insights about the entropic cost for peptide folding, and as such challenges the understanding of how likely it is that a particular peptide sequence would fold into the required conformation for binding.

Our strategy is to develop a reliable physics-based approach for predicting structures and relative energetics of protein receptor – peptide complexes that are guided by low information content NMR data, such as chemical shift data. Chemical shift data is particularly attractive for this application because it is (i) prerequisite for further NMR studies to provide higher information content data, and (ii) obtained by either solution state or solid state NMR studies. Backbone chemical shifts can be determined by solution state NMR for systems as large at 60 – 80 kDa^44^, and potentially for even large complexes using solid-state NMR. For weakly binding systems these data may need to be supplemented with the strongest backbone / backbone, backbone / methyl, backbone / aromatic, or methyl / methyl NOEs, that can generally be obtained even for modestly large protein-peptide complexes. Methods like HADDOCK already use this general approach^45,46^, addressing the challenge in interpreting the ambiguity in the CSP data, and are successful in predicting docking between folded proteins guided by such data. The key advantage of HADDOCK is speed since it is based on docking and heuristic scoring functions. However, the folding-upon-binding nature of protein-peptide complexes makes them particularly challenging systems for docking predictions with all available methods, including HADOCK, and especially difficult when the receptor can accommodate peptides in different binding modes and different peptide conformations (e.g. helices and strands as observed for ET-peptide complexes). The MELD+NMR approach uses simulations and statistical mechanics to identify low free energy states and is thus suitable to account for the entropic component of folding upon binding. The main advantage of such an approach is the production of models that agree with both a physical model and experimental data, providing valuable biophysical parameters such as relative free energies of binding.

In this study, we also observe that AlphaFold can successfully predict structures of these ET - peptide complexes. Similar applications of AlphaFold for peptide docking have also been recently reported by other groups^35^. However, the deep-learning based AlphaFold models do not provide information about relative binding affinities, which is a natural product of MELD binding simulations. In the case of the ET – JMJD6 complex, AlphaFold returns multiple models with different binding poses. In this sense, NMR-guided MELD+NMR and data-independent AlphaFold calculations provide both validating and complementary information for accurately modeling receptor-peptide complexes.

Here we demonstrate a successful approach for accurate modeling of protein-peptide complexes combining NMR backbone chemical shift data and MELD simulations. The goals of the blind study are multiple: (i) determine if the method is successful, (ii) assess its sensitivity to sequence and conformation, (iii) determine the amount of data needed for confident determination of the structures, and (iv) assess if MELD+NMR simulations can provide reliable relative free energies of binding. We observe that for the tight (K_d_ < 1 μM) and moderately tight (∼ 10 - 100 μM) ET – peptide complexes, MELD+NMR can reliably predict bound-state conformations of the peptide and relative binding free energies using only backbone chemical shift data (for both the protein receptor and for the bound peptide). Significantly, the method is successful in blind binding studies involving very different binding modes; e.g., the TP peptide and NSD3 peptide both bind ET as beta hairpins but bind in “flipped” orientations (see Fig. 1). In the best cases of the TP - ET, CHD4 - ET, and BRG1 -ET complexes, only CSP data on the receptor (i.e. ET) side of the complex was sufficient for an accurate NMR-guided MELD docking; however for the NSD3 - ET and LANA - ET complexes accurate modeling also required backbone chemical shifts for the bound peptide, which are used to define backbone dihedral restraints with TALOS. The MELD+NMR docking simulations also provide accurate relative free energies of binding for TP versus NSD3, in excellent agreement with experimental measurements.

In binding to peptides, the ET domain exhibits CSPs throughout the domain structure, reflecting an allosteric conformational change that is propagated across most of the domain^7^. The biological significance of these structural changes resulting from partner binding is not yet understood. As expected, the unstructured N-terminal regions of ET, which remains unstructured in the complex, does not exhibit CSPs due to peptide binding and/or allosteric changes^7^. The CSP data in Figure S1 shows insignificant chemical shift changes in this N-terminal unstructured region of ET, not involved in binding, where Δδ_(N,H)_ chemical shift changes between apo and peptide-bound forms are all < 0.02 ppm. For peptide docking, we set the threshold CSP as > 0.25 ppm - a threshold value that is enough to capture the highest perturbations, while bearing in mind that smaller Δδ_(N,H)_ CSPs (0.02 to 0.25 ppm) are also significant. As seen in Fig. 2 and Fig S1, with this CSP threshold we identify ∼ 20 residues as potentially involved in peptide binding – but, MELD is able to correctly identify that only a subset of these contacts are present in the peptide binding epitope of ET, with the rest of these CSPs corresponding to conformational changes of ET that accompany peptide binding. It is more difficult to obtain epitope-specific CSPs for weak binders, such as JMJD6 (see bottom panel in Fig. S1), where the low population of the bound state yields smaller changes in chemical shifts than observed for NSD3 or TP, precluding the use of the CSP data due to JMJD6 binding. Assuming competitive binding of two peptides for a common (or similar) binding site, transferring CSP data from strong binders for modeling complexes of weaker binders is thus a promising strategy for modeling their binding modes.

Using the published experimental chemical shift data for the weakly-binding JMJD6-BRD3 ET complex (K_d_ ∼ 160 µM^18^) in MELD-NMR docking simulations results in a binding pose similar to the experimental structure, where the binding mode is shifted by ∼6 Å RMSD relative to the native pose. Similar results were obtained using the BRD4 ET domain in the MELD-NMR docking calculations, and for models of this complex returned by AlphaFold (Fig. S5). We also carried out explicit solvent molecular dynamics simulations starting from the experimental NMR structure (without restraints). These simulations also exhibited shifting of the peptide out of the hydrophobic cleft of ET after 100 ns of simulation (Fig. S5). While the experimental structure shows a tryptophan residue of the JMJD6 buried in the hydrophobic cleft of the highly homologous BRD4 ET domain, this is not observed in the MELD-NMR, AlphaFold, or explicit solvent simulations starting from the experimental structure. These results suggest that the displaced pose observed in both MELD-NMR and AlphaFold might represent a true conformational state that is not modeled by the experimental NMR structure analysis. This discrepancy could arise from well-known challenges in interpretation of ensemble-averaged NMR data. Simulations provide the structure with the most weight in the Boltzmann ensemble. NOESY experiments yield peaks in the spectra when two atoms are close in space. The intensity of the peaks is an ensemble average over the experimental observable, which rapidly decays with increasing distance between the two atoms as <1/r^6^>^47,48^. In practical terms, short distances between two atoms have a stronger weight in conventional NMR structure determination protocols than those at a longer distance. A structure exhibiting a long distance between two atoms 70% of the time and a close distance 30% of the time, may be interpreted by conventional modeling methods as a short distance in one single (or dominant) conformation present in the sample. We believe, this could be of special importance in the case of weak peptide binders, such as JMJD6. If multiple binding modes are present, some of the NOEs observed may arise from a minor population with short interproton distances. Satisfying these NOE-based distance restraints could then overweight the representation of these structures in the final ensemble of prediction models. This is a general challenge for docking studies using NOE data. It provides a good illustration of why NMR-guided MELD+NMR protocols using exclusively chemical shift data may be even better suited for modeling the dominant poses of moderately-tight peptide – protein complexes than conventional methods using such ensemble-averaged NOE data.

In developing MELD+NMR we carefully considered the accessibly of experimental NMR data needed for successful modeling of complexes. The method assumes backbone ^15^N-^1^H CSPs for the receptor protein, which can generally be obtained for systems of up to 60 – 80 kDa. For studies of a series of peptides binding to a common receptor, it may often be sufficient to use the CSP data from one complex in modeling a second complex with a peptide known to complete for binding with the peptide of the first complex, as shown here for the TP-ET and NSD3-ET complexes. In some cases, it is also helpful to have backbone chemical shift data for the bound peptide for generating backbone dihedral angle restraints, requiring the production of isotope-enriched peptides. This is facilitated by several recently described peptide-protein fusion systems for high-level production of isotope-enriched peptides^7^. Although not the focus of the present study, other types of NMR such as residue dipolar couplings (RDCs) could be valuable to orient the peptide with respect the protein secondary structure elements^49^, albeit with the same caveats for the effects of dynamic averaging in weaker binding complexes discussed above for chemical shift and NOE data. The nature of such data requires new restraint types and replica exchange optimizations in MELD, which are planned developments for future work.

The eruption of Machine Learning into the field of structural biology has many exciting prospects. We took advantage of the AlphaFold pipeline as an orthogonal computational approach. Although AlphaFold was not originally designed for the purpose of peptide binding, a recent report shows that a tethered protein-peptide method using AlphaFold, similar to the approach used in this work, is about ∼40% accurate in peptide-protein docking benchmarks^35^. The AlphaFold predictions for three of the six systems analyzed in detail here were remarkably good matches to the MELD+NMR models. In particular, both provide accurate models of the TP-ET complex, and for the NSD3-ET complex both methods yielded a model where the structured domain was a hairpin, and the unstructured termini formed a helix (with the same orientation). Such agreement highlights a tendency from both the physics model and the deep-learning algorithm towards helical states for this generally-dynamic region of the sequence (see Fig. 4). For the JMJD6 peptide, both methods modeled a helix as the bound-state peptide conformation, consistent with the experimental structure, and both predicted a binding mode displaced with respect the experimentally - reported binding site (Fig. S2). For the three other systems studied, AlphaFold can capture the native states but the relative orientation between the receptor and peptide residues is shifted by one or two residues compared to the experimental structures (Fig. S6). MELD_NMR, however, provided models for these complexes in excellent agreement with the experimental structures. Overall, we take these predictions with optimism that deep-learning approaches such as AlphaFold can be used to filter and identify where and how peptides are likely to interact with a protein receptor. However, as they are predictions, they need to be compared with at least some experimental data before building studies based on these models. The MELD+NMR approach used here is more computationally expensive at run time but produces models that are already compatible with experimental NMR data. MELD+NMR accounts for peptide conformational entropic preferences through sampling, and can be exploited for virtual competitive binding studies to assess binding affinities. Powerful strategies will combine both methods in assessing peptide ligand candidates, with structural filtering with AlphaFold to select peptides worth exploring with experimental NMR studies followed by MELD+NMR to produce experimentally-refined models and to rank order binding preferences.

## CONCLUSION

The novel MELD+NMR protocol produces valuable insights and predictions for the problem of polypeptide folding upon binding, a challenging area of computational modeling. We have shown reliable predictions for tight and moderately tight (K_d_ < 10 - 100 μM) peptide - protein complexes using only backbone chemical shift data, for both the receptor and the bound peptide, together with simulations. However, a few limitations are apparent. For weaker binders the current protocols struggle to find the correct conformation without at least some NOE-based distance restraints. This is attributable, at least in part, to conformational averaging in weaker complexes, which confounds the interpretation of NMR data. Future work will aim to improve performance in cases of weak binders where several conformations might be contributing to the binding affinity. This tandem experimental/computational approach can also be useful for rational peptide design, where a large number of peptide designs directed to common receptor can be screened for potential complex formation with AlphaFold and then tested using MELD-NMR.

## Supporting information

Supplemental tables and results

## ASSOCIATED CONTENT

### Supporting Information

Supplementary Results and Tables S1-9 are provided as a PDF. Supplementary Figures S1-6 are provided as a PDF.

## AUTHOR INFORMATION

### Author Contributions

AM, GVTS, MJR, GTM, and AP designed and supervised the research. AM, GVTS, and AP carried out computational methods development, computational modeling calculations, and model analysis. JH and LM carried out experimental studies. The manuscript was written with contributions from all authors. All authors have given approval to the final version of the manuscript.

### Notes

GTM is a founder of Nexomics Biosciences, Inc. This affiliation is not a competing interest with respect to this study. The remaining authors declare no competing interests.

## ACKNOWLEDGMENT

We thank Dr. Theresa Ramelot for useful suggestions in the course of this study.

## ABBREVIATIONS

BET: Bromo and extraterminal domain protein
BRD3 and BRD4: Specific BET proteins
CSP: Chemical Shift Perturbation
ET: Extraterminal domain (residues 554 to 640) of human BRD3
MELD: Modeling Employing Limited Data
NSD3: Peptide fragment (residues 148 – 184) of the human Nuclear Receptor Binding SET Domain Protein 3
TP: Peptide corresponding to 22 C-terminal residues of murine ET

## SYNOPSIS TOC

The Extraterminal domain of BET proteins accommodates binding of different peptide sequences along different binding modes. We combine computational methods designed to handle ambiguous and sparse contact data together with NMR chemical shift data to determine structures of these complexes in excellent agreement with experimental structures determined using extensive NMR data sets.

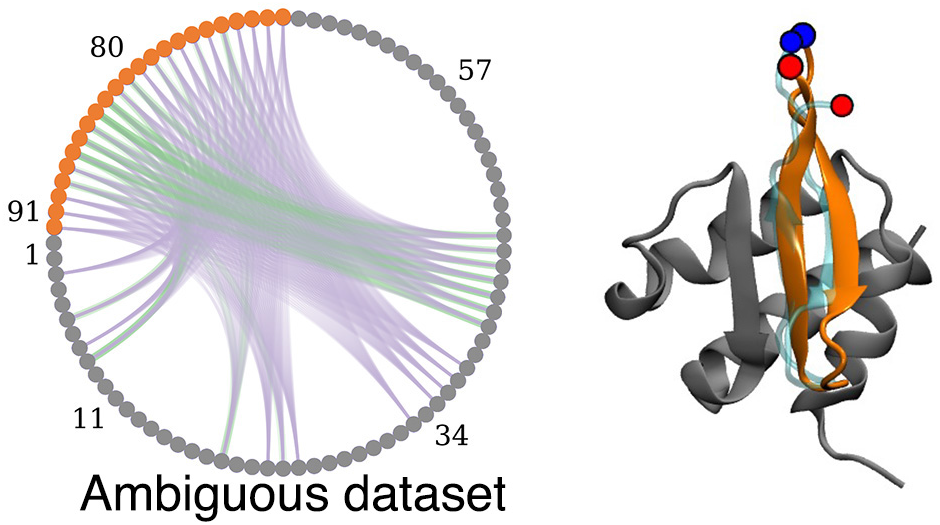

