## Supplemental tables and results for "Structure determination of protein-peptide complexes from NMR chemical shift data using MELD"

### Supplementary Results:

Transferring ET-TP CSP data to ET-NSD3: For NSD3 peptide simulations using the CSP+TALOS data guided the system to the native state (IRMSD=2.0, ILRMSD=2.9, $f_{nat}$ (fraction of native contacts) =72.9%). Repeating the simulation using CSP experimental data from the ET-TP system yielded predictions of similar accuracy (IRMSD=2.5 Å, ILRMSD=2.9Å, and $f_{nat}$=79%). In both simulations the top cluster had a population of 49%.

Modeling CSP Data: We have taken two approaches to modeling the data. In the first one, we create the combinatorics for all peptide residues with *active* protein residues. For the second one we only create combinatorics between hydrophobic pairs in the two sets. Simulations yielded results of similar quality for all peptides. In the case of BRD3-BRG1 only the hydrophobic pairing selections successfully predicted the experimental structure as the top cluster.

Additional Systems: We applied our MELD-NMR pipeline to study three additional peptides that have been experimentally characterized: BRD4-LANA, BRD3-CHD4 and BRD3-BRG1 (PDB IDs are 2ND0, 6BGG and 6BGH respectively, see Table 1). These three peptides bind as a small antiparallel single β-strand. Given the homology and structural similarity between the ET domains of BRD3 and BRD4, we simulated the BRD3-LANA binding for consistency with the other systems studied in this work.

**CSP dataset:** We transferred the *CSP* dataset of the BRD3-TP system as described in the methods section to study these three systems. With this information, MELD predicts the native complex structure correctly for only BRD3-CHD4 system (40% population excluding termini, see Fig. S4). The case of BRD3-BRG1 is interesting since it contains a symmetric binding motif and MELD simulations could not differentiate between an antiparallel or parallel β-strand orientation – with the top cluster incorrectly favoring the parallel orientation when using all possible pairings between residues selected as *active* and peptide residues. Using only the hydrophobic residues of the peptide and the protein result in native antiparallel binding mode with top cluster population around 31%. For BRD3-LANA, MELD could not find the peptide's experimental binding site (Fig. S4).

**CSP + TALOS dataset:** Like JMJD6, the LANA peptide is a weak binder (with binding affinity 635 μM), and indeed TALOS predictions based on chemical shift shows backbones of this peptide to be dynamics, offering no guiding power for the simulations (see Table S5). The chemical shift lists for BRD3-CHD4 (BMRB code 30367) and BRD3-

BRG1 (BMRB code 30368) systems lacked information about C, N atoms which are essential for TALOS to predict backbone dihedral data. Thus, for these three systems we used simulated backbone phi and psi data back calculated from the experimental structure. We enforced phi/psi ranges for the 3 complexes as described in the methods section with standard deviation of 40 degrees for each dihedral (see Table S4, S5, and S6). CHD4 and BRG1 continue to be in good agreement with experiments, with an increased population of top cluster. The MELD prediction for BRD3-LANA complex is partially correct, with the C-terminal floppy region folding back to form a hairpin like structure (Fig. S4).

**CSP + TALOS + NOE dataset:** Our third type of simulations add the three strongest NOEs for each system. We left out BRD3-CHD4 and BRD3-BRG1 as they were already successful and the BMRB entry contained no NOE peak lists. The BMRB entry for BRD4-LANA (BMRB code 26042) did not contain the NOE peak list either, hence we took the three shortest backbone H-H distances back-calculated from the experimental structures (44H - 75H, 42H - 77H, and 44H - 76H). We enforced them in the MELD simulations as described in the methods section. This resulted in higher accuracy predictions, with the top cluster (with population of 48%) now representing the experimental binding mode (Fig. S4).

We complemented our study with AlphaFold predictions (see methods), finding good agreement with experiments for all three systems (Fig. S4). Upon comparing these predictions with MELD successful predictions, we realized the lRMSD values are similar for both MELD and AlphaFold2 (AF2) predicted structures, whereas the $f_{nat}$ values are significantly lower for AF2 predicted structures compared to MELD predictions. Upon further inspection of these systems, the registry between the residue pairing in the receptor and peptide strand is shifted by two residues with respect to the experimental structures (Fig. S6).

**Supplementary Tables**:

**Table S1**. List of backbone dihedral angle restraint ranges used in MELD for the TP peptide These dihedral angle restraints were calculated from peptide chemical shift using TALOS+. Peptide residue numbering follow the numbering in MELD simulations (see Table 1). We only used dihedral angle restraint ranges that are categorized as *strong* in the TALOS+ output file.

| Residue number | Residue name | PHI-min | PHI-max | PSI-min | PSI-max |
|---|---|---|---|---|---|
| 72 | T | -167.8 | -64.6 | 109.7 | 173.8 |
| 73 | W | -151.1 | -73.6 | 93.6 | 194.8 |
| 74 | R | -155.6 | -103.6 | 133.5 | 181.6 |
| 75 | V | -151.0 | -64.6 | 105.6 | 145.6 |
| 76 | Q | -162.0 | -54.6 | 111.2 | 162.2 |
| 77 | R | -119.1 | -51.4 | 119.5 | 159.5 |
| 78 | S | -147.9 | -59.8 | 96.7 | 199.0 |
| 80 | N | -156.2 | -79.4 | 35.6 | 175.6 |
| 83 | K | -178.8 | -44.3 | 123.6 | 178.4 |
| 84 | I | -152.3 | -112.3 | 131.0 | 171.0 |
| 85 | R | -164.2 | -98.2 | 105.1 | 165.8 |
| 86 | L | -134.1 | -94.1 | 109.6 | 164.6 |
| 87 | T | -139.5 | -97.9 | 111.8 | 160.4 |
| 88 | R | -126.2 | -79.0 | 98.8 | 148.2 |
| 89 | E | -138.1 | -55.4 | 92.5 | 157.0 |
| 90 | A | -140.3 | -5.0 | 119.0 | 169.2 |

**Table S2**. List of backbone dihedral angle restraint ranges used in MELD simulations for NSD3 peptide. These dihedral angle restraints were calculated from peptide chemical shift using TALOS+. Peptide residues number correspond to the residue numbering in MELD simulations (Table 1). We only used dihedral angle restraint ranges that are categorized as *strong* in the TALOS+ output file.

| Residue number | Residue name | PHI-min | PHI-max | PSI-min | PSI-max |
|---|---|---|---|---|---|
| 71 | I | -137.9 | -57.9 | 79.4 | 179.4 |
| 72 | K | -175.0 | -95.0 | 109.3 | 209.3 |
| 73 | L | -176.0 | -96.0 | 90.6 | 190.6 |
| 74 | K | -154.4 | -74.4 | 74.8 | 174.8 |
| 75 | I | -148.9 | -68.9 | 75.7 | 175.7 |
| 76 | T | -159.0 | -79.0 | 70.4 | 170.4 |
| 77 | K | -150.9 | -70.9 | 71.6 | 171.6 |
| 78 | T | -163.5 | -83.5 | 85.6 | 185.6 |
| 79 | I | -150.8 | -70.8 | 73.4 | 173.4 |
| 80 | Q | -152.7 | -72.7 | 90.7 | 190.7 |
| 84 | E | -131.5 | -51.5 | 76.0 | 176.0 |
| 85 | L | -151.9 | -71.9 | 84.5 | 184.5 |
| 86 | F | -164.8 | -84.8 | 104.4 | 204.4 |
| 87 | E | -183.9 | -103.9 | 101.0 | 201.0 |
| 88 | S | -167.0 | -87.0 | 84.6 | 184.6 |
| 89 | S | -161.6 | -81.6 | 100.8 | 200.8 |

Table S3. List of backbone dihedral angle restraint ranges used in MELD simulations for JMJD6 peptide. Dihedral angle restraints for each residue in the peptide are calculated using based on the experimentally solved NMR ensemble (6BNH) using MDTraj. In MELD simulations, we used dihedral angle restraint ranges only for the residues that a have narrow distribution of dihedral values for 20 NMR structures.

| Residue number | Residue name | PHI-min | PHI-max | PSI-min | PSI-max |
|---|---|---|---|---|---|
| 70 | W | -181.1 | -101.1 | -4.0 | 76.0 |
| 71 | T | -118.4 | -38.4 | -207.4 | -127.4 |
| 72 | L | -82.5 | -2.5 | -77.5 | 2.5 |
| 73 | E | -95.7 | -15.7 | -110.4 | -30.4 |
| 74 | R | -104.9 | -24.9 | -97.1 | -17.1 |
| 75 | L | -104.8 | -24.8 | -103.4 | -23.4 |
| 76 | K | -102.2 | -22.2 | -77.1 | 2.9 |
| 77 | R | -122.0 | -42.0 | -53.1 | 26.9 |
| 78 | K | -136.7 | -96.7 | -54.9 | 25.1 |
| 79 | Y | -129.3 | -49.3 | -97.1 | -17.1 |
| 80 | R | -116.1 | -36.1 | -63.0 | 17.0 |

Table S4. List of backbone dihedral angle restraint ranges used in MELD simulations for LANA peptide. Dihedral angles restraints for each residue in the peptide are calculated using based on the experimentally solved NMR ensemble (2ND0) using MDTraj. In MELD simulations, we used dihedral angle restraint ranges only for the residues that have narrow distribution of dihedrals for 20 NMR structures.

| Residue number | Residue name | PHI-min | PHI-max | PSI-min | PSI-max |
|---|---|---|---|---|---|
| 74 | I | -116.8 | -36.8 | 80.0 | 160.0 |
| 75 | V | -140.8 | -60.8 | 109.8 | 189.8 |
| 76 | K | -174.1 | -94.1 | 87.8 | 167.8 |
| 77 | F | -180.6 | -100.6 | 111.6 | 191.6 |
| 78 | K | -137.5 | -57.5 | 91.5 | 171.5 |

Table S5. List of backbone dihedral angle restraint ranges used in MELD simulations for BRG1 peptide. Dihedral angle restraints for each residue in the peptide are calculated using based on the experimentally solved NMR ensemble (6BGH) using MDTraj. In MELD simulations, we used dihedral angle restraint ranges only for those residues that have narrow distribution of dihedral values for 20 NMR structures.

| Residue number | Residue name | PHI-min | PHI-max | PSI-min | PSI-max |
|---|---|---|---|---|---|
| 71 | V | -136.6 | -56.6 | - | - |
| 72 | K | - | - | 76.7 | 156.7 |
| 73 | V | -124.3 | -44.3 | - | - |
| 74 | K | -174.2 | -94.2 | 42.1 | 122.1 |
| 75 | I | -120.0 | -40.0 | - | - |
| 76 | K | - | - | 48.8 | 128.8 |
| 77 | L | -117.5 | -27.5 | - | - |

Table S6. List of backbone dihedral angle restraint ranges used in MELD simulations for CHD4 peptide. Dihedral angle restraints for each residue in the peptide are calculated using based on the experimentally solved NMR ensemble (6BGG) using MDTraj. In MELD simulations, we used dihedral angle restraint ranges only for those residues that have narrow distribution of dihedral values for 20 NMR structures.

| Residue number | Residue name | PHI-min | PHI-max | PSI-min | PSI-max |
|---|---|---|---|---|---|
| 70 | V | -171.0 | -91.0 | 97.6 | 177.6 |
| 72 | P | - | - | 98.6 | 218.6 |
| 73 | L | -111.4 | -31.4 | 55.2 | 135.2 |
| 74 | K | -180.0 | -100.0 | - | - |
| 75 | I | -194.5 | -114.5 | 95.5 | 175.5 |
| 79 | G | -134.0 | -54.0 | - | - |
| 80 | F | -168.6 | -88.6 | - | - |

Table S7. List of predicted backbone dihedral angles from TALOS-N for bound JMJD6 based on backbone chemical shift data.

| RESID | RESNAME | PHI | PSI | DPHI | DPSI | DIST | S2 | COUNT | CS_COUNT | CLASS |
|---|---|---|---|---|---|---|---|---|---|---|
| 69 | K | - | - | 0 | 0 | 0 | 0 | 0 | 6 | None |
| 70 | W | -67.0 | 136.59 | 12.87 | 6.83 | 1.403 | 0.27 | 6 | 10 | Dyn |
| 71 | T | -104.2 | 145.97 | 31.86 | 18.85 | 0.942 | 0.37 | 25 | 11 | Dyn |
| 72 | L | -64.76 | -30.81 | 5.85 | 7.20 | 0.838 | 0.5 | 25 | 12 | Dyn |
| 73 | E | -68.50 | -21.25 | 6.82 | 7.79 | 0.810 | 0.635 | 25 | 12 | Strong |
| 74 | R | -74.68 | -20.76 | 7.98 | 8.75 | 0.719 | 0.615 | 25 | 12 | Strong |
| 75 | L | -70.98 | 139.46 | 8.27 | 7.07 | 0.923 | 0.558 | 9 | 12 | Dyn |
| 76 | K | -75.95 | -20.43 | 14.67 | 20.61 | 2.724 | 0.486 | 1 | 12 | Dyn |
| 77 | R | -69.14 | -33.55 | 17.70 | 16.91 | 0.000 | 0.431 | 25 | 12 | Dyn |
| 78 | K | -76.62 | -30.43 | 17.55 | 18.25 | -0.90 | 0.388 | 7 | 12 | Dyn |
| 79 | Y | -97.77 | 114.36 | 24.77 | 41.84 | 0.000 | 0.318 | 10 | 12 | Dyn |
| 80 | R | -75.4 | -26.96 | 18.85 | 19.40 | 0.000 | 0.258 | 10 | 12 | Dyn |
| 81 | N | - | - | 0 | 0 | 0 | 0 | 0 | 8 | None |

Table S8. List of predicted backbone dihedral angles from TALOS-N for bound LANA based on backbone chemical shift data.

| RESID | RESNAME | PHI | PSI | DPHI | DPSI | DIST | S2 | COUNT | CS_COUNT | CLASS |
|---|---|---|---|---|---|---|---|---|---|---|
| 69 | N | - | - | 0 | 0 | 0 | 0 | 0 | 9 | None |
| 70 | L | -69.82 | 137.21 | 9.93 | 9.77 | 0.507 | 0.39 | 25 | 14 | Dyn |
| 71 | Q | -68.42 | 136.06 | 6.11 | 8.72 | 0.442 | 0.60 | 25 | 15 | Warn |
| 72 | S | -72.85 | 144.04 | 9.67 | 9.36 | 0.385 | 0.61 | 10 | 15 | Warn |
| 73 | S | -70.22 | 143.27 | 6.99 | 13.73 | 0.387 | 0.54 | 9 | 15 | Dyn |
| 74 | I | -78.00 | 135.29 | 15.37 | 14.47 | 0.446 | 0.39 | 10 | 15 | Dyn |
| 75 | V | -77.80 | 133.68 | 15.11 | 14.96 | 0.438 | 0.39 | 25 | 15 | Dyn |
| 76 | K | -77.92 | 126.97 | 7.19 | 16.86 | 0.475 | 0.45 | 25 | 15 | Dyn |
| 77 | F | -68.58 | -25.88 | 12.28 | 14.11 | 0.639 | 0.67 | 8 | 14 | Warn |
| 78 | K | -64.61 | -26.75 | 6.19 | 8.42 | 0.665 | 0.75 | 25 | 14 | Strong |
| 79 | K | -69.35 | 134.23 | 9.82 | 10.76 | 3.211 | 0.73 | 9 | 12 | Warn |
| 80 | P | -68.01 | 146.20 | 6.09 | 12.90 | 1.167 | 0.59 | 10 | 13 | Dyn |
| 81 | L | -74.09 | 134.85 | 6.85 | 11.54 | 0.439 | 0.50 | 25 | 11 | Dyn |
| 82 | P | -67.13 | 144.88 | 6.70 | 8.39 | 0.339 | 0.44 | 25 | 13 | Dyn |
| 83 | L | -73.4 | 132.37 | 11.88 | 12.06 | 0.310 | 0.46 | 10 | 13 | Dyn |
| 84 | T | -78.1 | 144.66 | 12.5 | 14.50 | 0.356 | 0.45 | 10 | 15 | Dyn |
| 85 | Q | -69.83 | 125.78 | 7.11 | 11.39 | 0.443 | 0.43 | 25 | 13 | Dyn |
| 86 | P | -63.84 | 148.78 | 8.6 | 8.9 | 0.580 | 0.41 | 25 | 12 | Dyn |
| 87 | G | - | - | 0 | 0 | 0 | 0 | 0 | 7 | None |

Table S9. Experimental NOEs for the ET-JMJD6 system, chosen from NOE peak list in four different categories: protein backbone / peptide backbone, protein methyl sidechain/ peptide backbone, protein backbone / peptide methyl sidechain, and protein methyl sidechain / peptide methyl sidechain. In the second row, we further included peaks coming from single Trp70 residue in peptide. Residue numbering corresponds to the residue range in MELD (Table 1).

| | Backbone-Backbone | Sidechain-Backbone | Backbone - Sidechain | Methyl - Methyl |
|---|---|---|---|---|
| Experimental NOE with ILVA methyl | Not present | 20 HD21-75HA | 13 HA - 75 HD11<br>9 HA - 75 HD11 | 12 HD21 - 75 HG21<br>20 HD11 - 75 HD21<br>23 HD11 - 75 HG21 |
| Experimental NOEs with ILV methyl and Trp70 | Not present | 20 HD21-75HA | 13 HA - 75 HD11<br>9 HA - 75 HD11<br>6 HA - 70 HH2 | 12 HD21 - 75 HG21<br>20 HD11 - 75 HD21<br>44 HD12 - 70 HH2 |
