## Supplemental figures for "Structure determination of protein-peptide complexes from NMR chemical shift data using MELD"

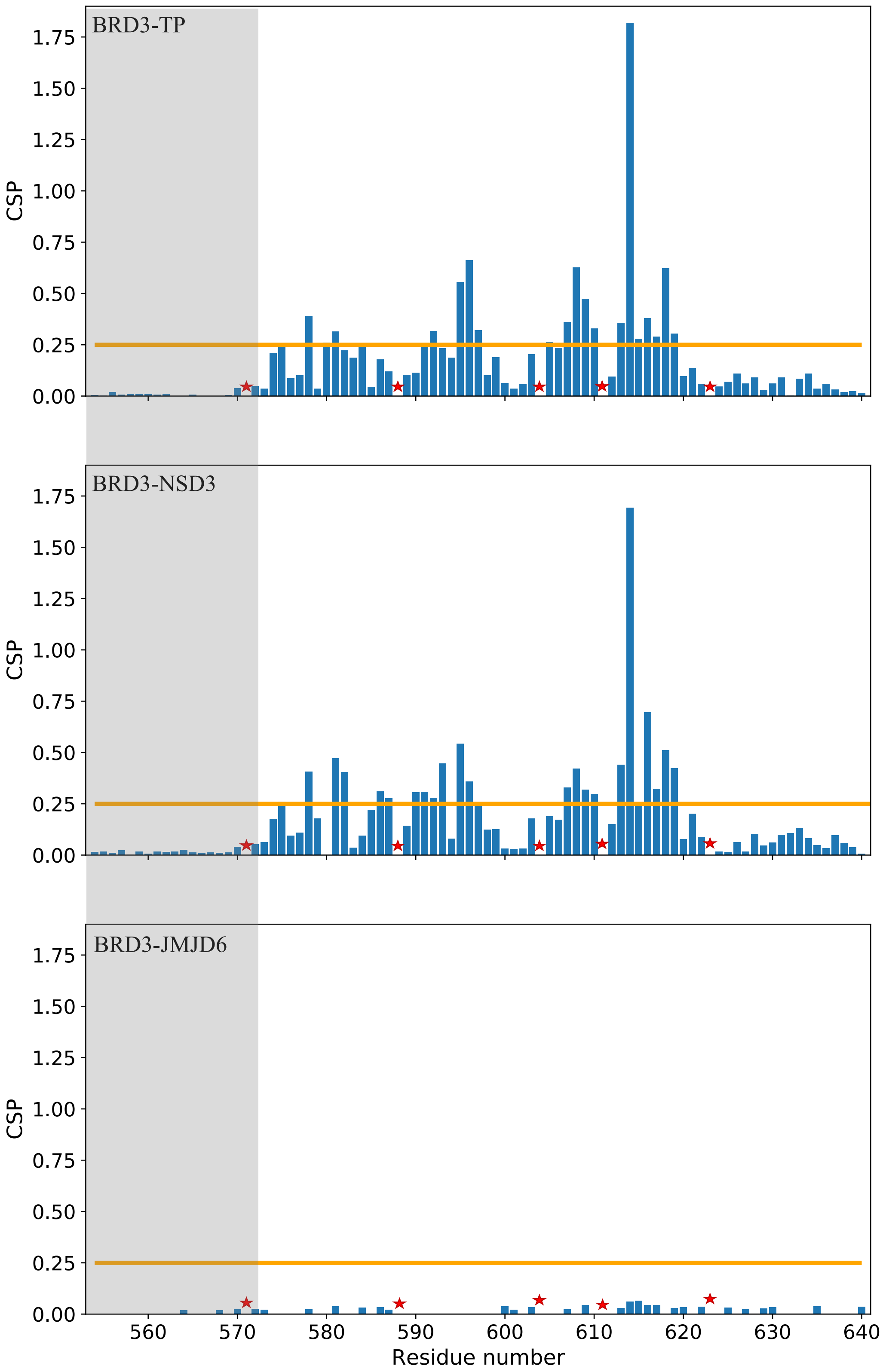


**Figure S1**. **CSP values for each residue along the ET domain represented as bar plots**. The orange line shows the threshold used to define *active residues* for the binding process. The shaded region corresponds to the floppy tail in the receptor which was excluded from simulations. The star symbols denote residues for which CSPs could not be measured (e.g. Pro residues). CSP measurement for both ET-TP and ET-NSD3 were performed at pH=7.0 with a peptide concentration 0.5 μM and a 1:1 ratio of protein to peptide.

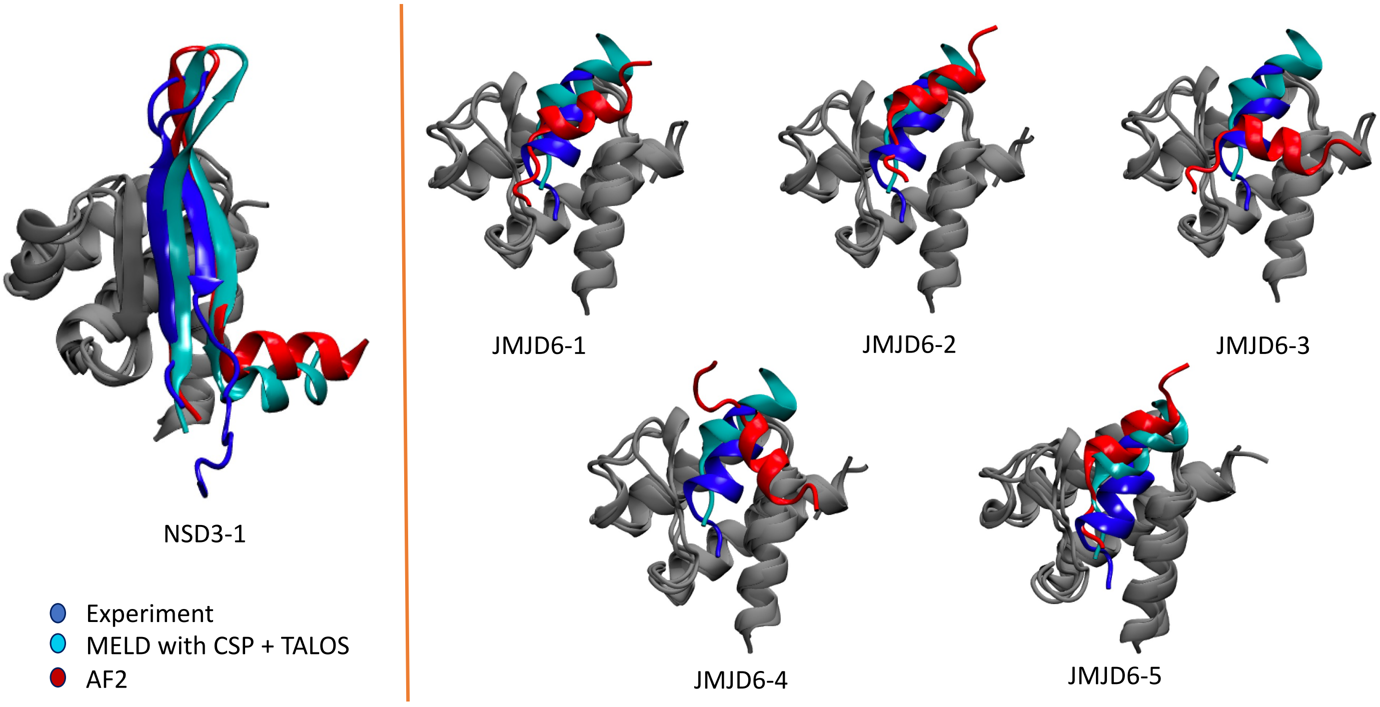


**Figure S2.** Superposition of the experimental structure, MELD prediction and AlphaFold's best prediction for the ET-NSD3 system (left panel). Superposition of the experimental structure, MELD prediction and top five AlphaFold's prediction for the ET-JMJD6 system (right panel).

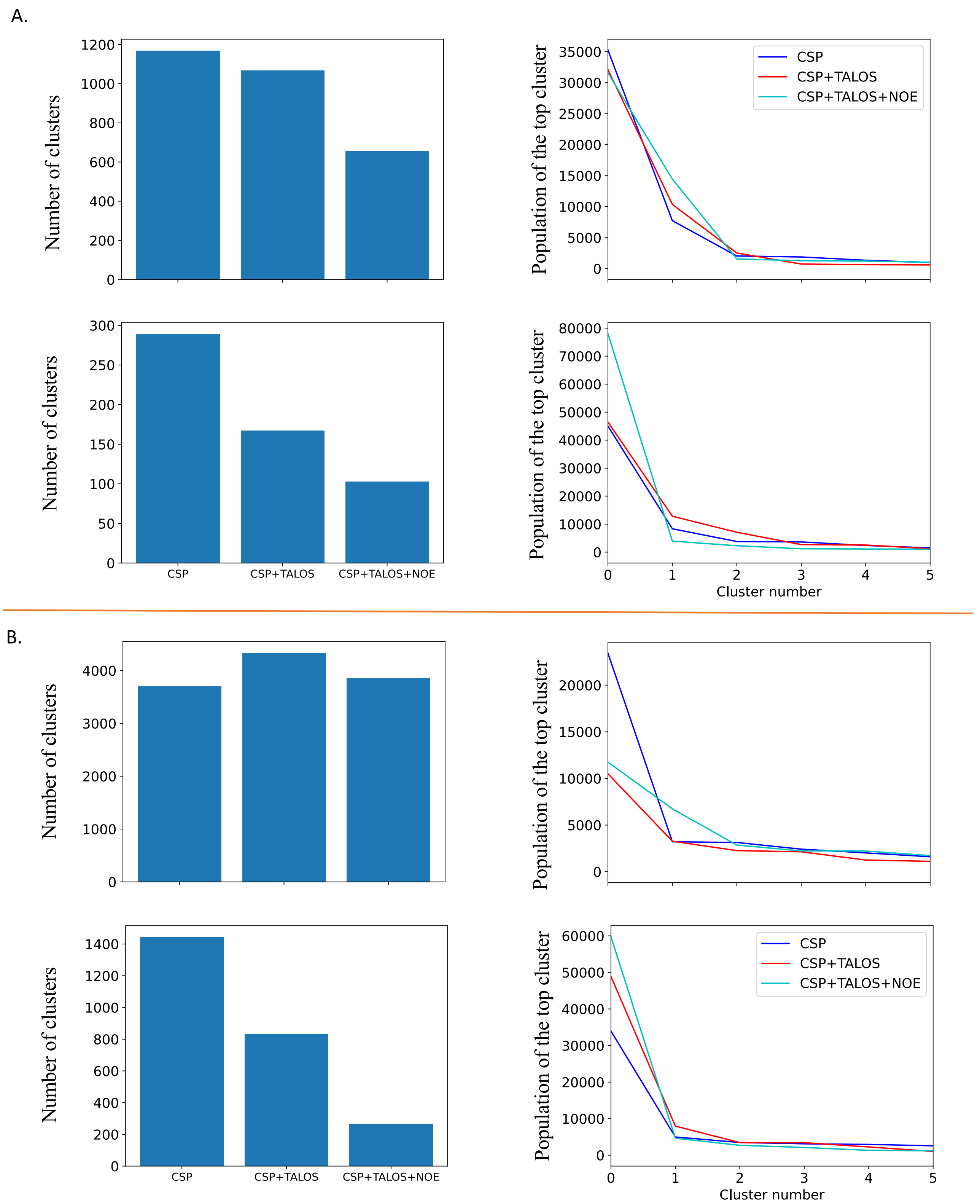


**Figure S3.** The amount of information used dictates the ability to sample multiple binding modes (left) and identify the native-like one as the highest population cluster (right). (A.) ET- TP and (B.) ET-NSD3(B.). The top rows of panel A and panel B correspond to clustering over the whole peptide, while the bottom rows exclude the floppy termini (3 residues for TP and 11 residues for NSD3).

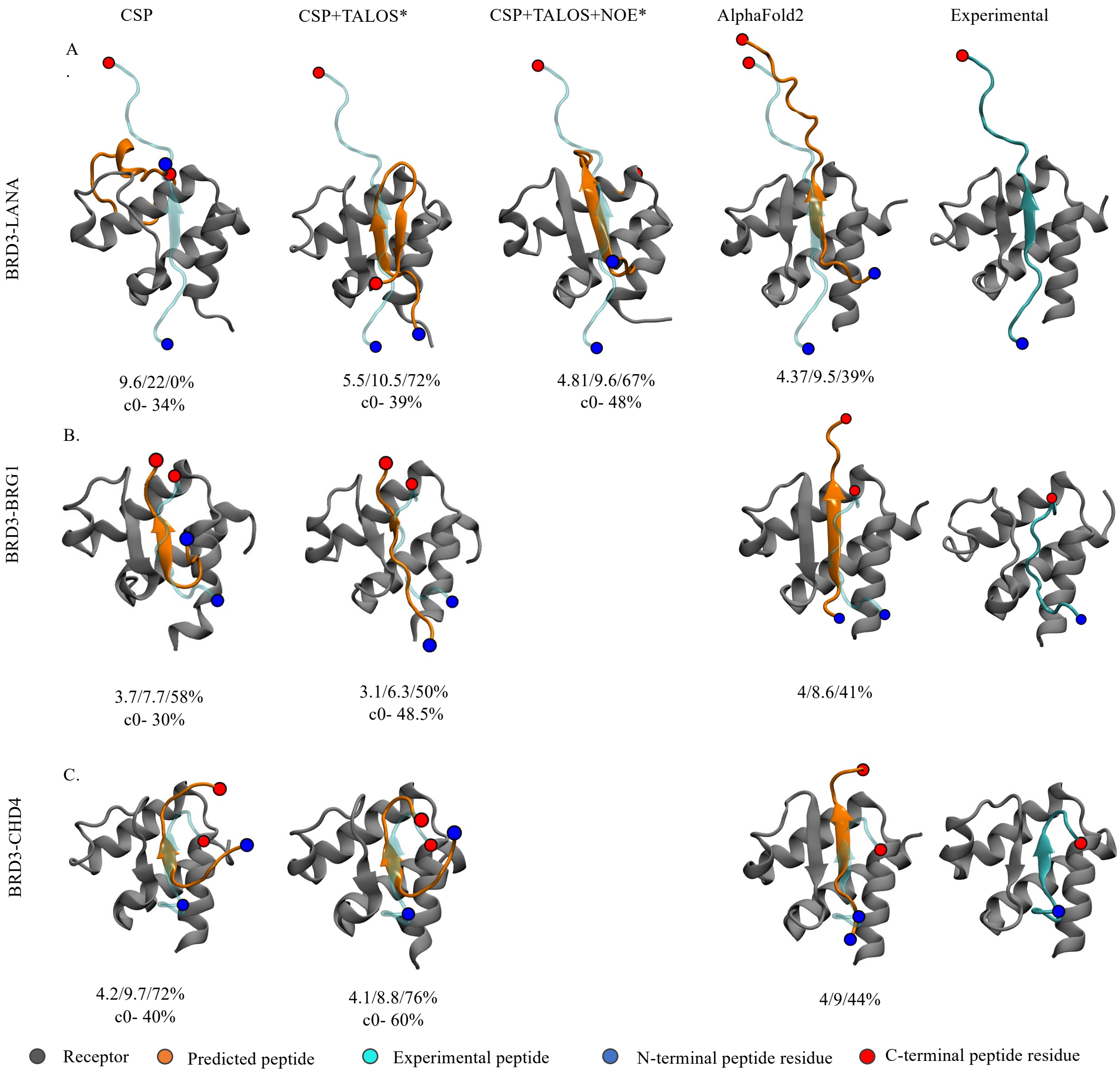


Figure S4. MELD prediction using different dataset (first three columns) for the extended study along with AlphaFold predictions (4$^{th}$ column) and experimental structure (5$^{th}$ column): (A.) ET-LANA, (B.) ET-BRG1, and (C.) ET-CHD4. The numbers stand for IRMSD/ILRMSD/$f_{nat}$ in the first row and the population of the top clusters in the second row.

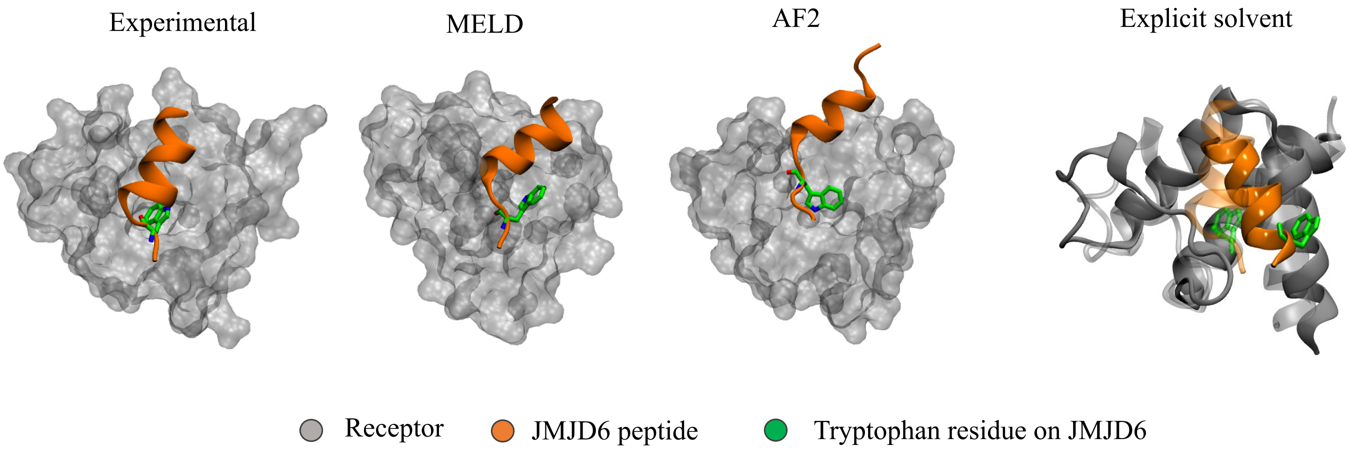


**Figure S5**. Orientation of tryptophan residues in experiment and simulations. Both implicit and explicit simulations as well as AlphaFold favor conformations in which the tryptophan is not buried in the ET cleft, favoring a change in the binding mode with respect to experiments.

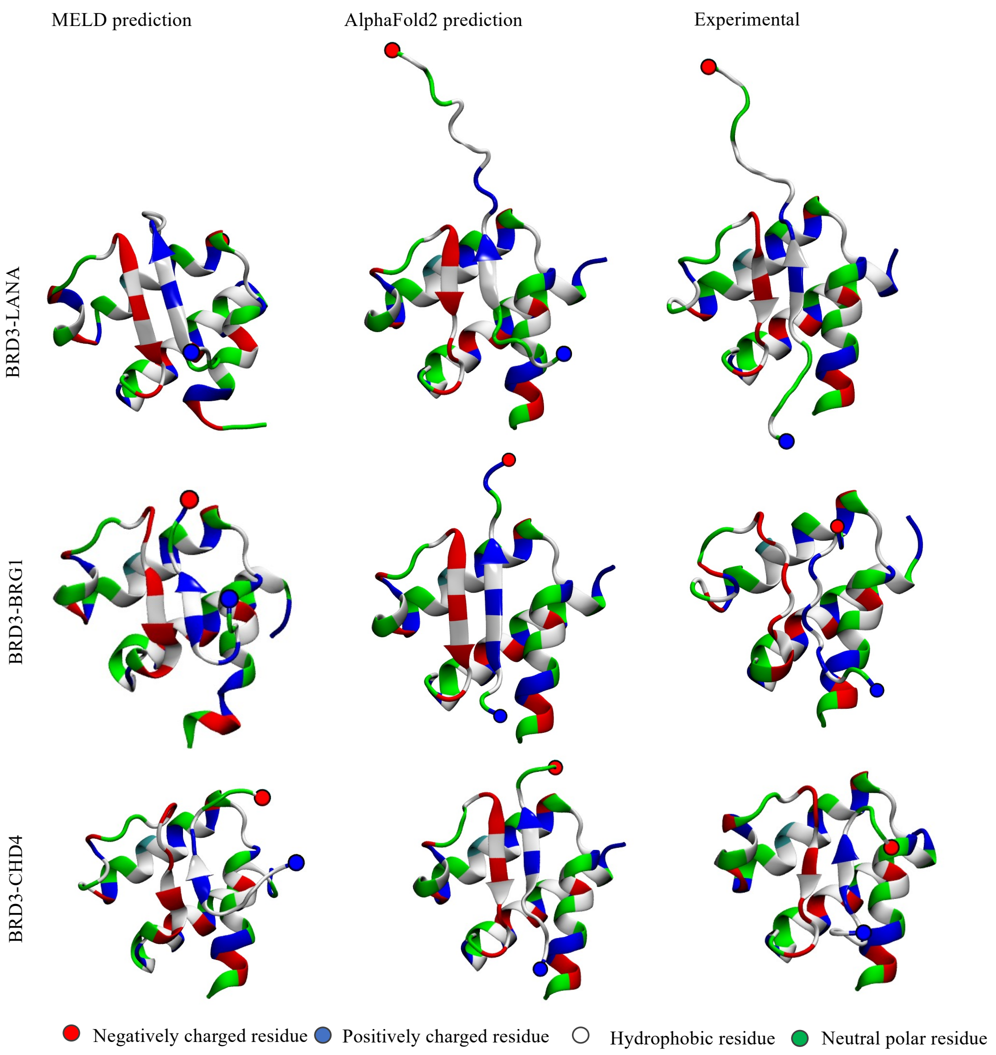


**Figure S6.** Residue type representation for the  top MELD prediction (left column), top AlpahFold prediction (middle column) and the experimental structure (right column).
